# Modeling prediction error improves power of transcriptome-wide association studies

**DOI:** 10.1101/108316

**Authors:** Kunal Bhutani, Abhishek Sarkar, Yongjin Park, Manolis Kellis, Nicholas J. Schork

## Abstract

Transcriptome-wide association studies (TWAS) test for associations between imputed gene expression levels and phenotypes in GWAS cohorts using models of transcriptional regulation learned from reference transcriptomes. However, current methods for TWAS only use point estimates of imputed expression and ignore uncertainty in the prediction. We develop a novel two-stage Bayesian regression method which incorporates uncertainty in imputed gene expression and achieves higher power to detect TWAS genes than existing TWAS methods as well as standard methods based on missing value and measurement error theory. We apply our method to GTEx whole blood transcriptomes and GWAS cohorts for seven diseases from the Wellcome Trust Case Control Consortium and find 45 TWAS genes, of which 17 do not overlap previously reported case-control GWAS or differential expression associations. Surprisingly, we replicate only 2 of 40 previously reported TWAS genes after accounting for uncertainty in the prediction.

## 2 Introduction

Thousands of loci associated with hundreds of complex diseases have been reported in the NHGRI catalog of genome-wide association studies [25, 50] (GWASs). However, most genome-wide significant loci are devoid of protein-coding alterations [18] and likely instead affect gene regulation by mechanisms such as transcriptional, post-transcriptional, or epigenetic regulation. Several studies have directly investigated the role of transcriptional regulation on complex diseases by jointly considering genotypes, expression, and phenotypes using Mendelian randomization [44, 3]. However, such studies require genetic, transcriptomic, and phenotypic data to be measured in all samples, which is still prohibitive at the scale of GWAS.

Recent large-scale efforts such as the Gene-Tissue Expression Project (GTEx) have generated reference transcriptomes across multiple human tissues[4]. These data have enabled transcriptome-wide association studies (TWAS), which use the reference expression data to build models of transcription regulation, impute gene expression into GWAS cohorts where expression is not measured, and directly test for association between predicted gene expression and phenotype[15, 16]. However, current methods are limited to using only point estimates of imputed expression, while ignoring the uncertainty in the predicted expression.

The impact of not incorporating uncertainty of imputed predictors on genetic association analysis has been previously studied in the context of imputed genotypes in GWAS. Although taking the best-guess genotype (posterior mode) is standard practice for GWAS, using posterior mean dosages increases power to detect associations [56, 1]. Standard methods from missing data theory such as multiple imputation have been applied in this setting yielding reductions in bias [42, 35]. More sophisticated (and computationally expensive) methods such as SNPTEST [33] can analytically integrate over the full posterior distribution of the imputed genotypes, further improving power.

Here, we develop a novel Bayesian method for modeling uncertainty in imputed expression and propagating this uncertainty through TWAS. We compare our method to existing methods for TWAS and standard methods from missing data and measurement error theory and show that our method increases power to detect genes associated with phenotype. We apply our methods to GWAS for seven diseases from the Wellcome Trust Case Control Consortium [9] and find 45 TWAS genes, replicating only 2 of 40 previously reported TWAS genes. We find 17 of the 45 genes have not yet been identified by GWAS or differential gene expression in case-control cohorts.

## 3 Results

### 3.1 Uncertainty in TWAS

The key insight enabling TWAS is that one can train models to predict gene expression from genotype in reference cohorts, and use these models to impute unobserved gene expression values in GWAS samples using the genotype information. Direct tests for association between gene expression and phenotype can be pursued, more directly identifying putative causal genes for the phenotype of interest. However, current methods only use a point estimate of the predicted gene expression in TWAS and ignore the uncertainty in the prediction. The uncertainty arises from two sources: not learning the correct model for transcriptional regulation (e.g., omitting trans-regulatory effects), which we do not consider here, and not correctly estimating the model parameters due to sample size, linkage disequilibrium, or biological and technical confounders.

Our main contribution is a novel two-stage Bayesian regression model which incorporates uncertainty for each SNP effect (BAY-TS). The key idea of BAY-TS is to use the posterior distribution of SNP effects from the first-stage regression of expression against genotype as the prior distribution on effects in the second stage regression of phenotype against expression (Methods). After performing the second stage regression, we compute a Bayes factor comparing the fitted model against a null model where gene expression has no effect on phenotype.

We compared our method against existing methods, which merely calculate association statistics using ordinary least squares. We compared two strategies: using the full elastic net model trained on the entire GTEx dataset (OLS-E), and using the mean value of imputed expression across 50 bootstrapped models (OLS-M). We note that existing methods for TWAS implement OLS-E only.

We then compared our method to multiple imputation (MI), a standard method for handling completely missing (i.e., unobserved) data such as gene expression in TWAS [30]. For each gene, we imputed 50 expression levels for each individual using the bootstrapped models described above and estimated 50 effect sizes. We then combined these effect sizes into a single association statistic for evaluation incorporating both the mean and the variance of the 50 estimates (Methods).

Finally, we compared our method to regression calibration (RC), a standard method from measurement error theory. Measurement error theory explicitly models the error in observations (here, imputed expression) and predicts the impact of not including the errors on statistical inference [14]. Briefly, not explicitly including error in the model leads to a violation of the model assumptions and therefore leads to bias in the estimated regression coefficients. Applying this theory to TWAS, we modeled each imputed expression value as the true expression value plus additive error. We estimated the distribution of the error as the variance in predicted expression across the 50 bootstrapped models. We then performed RC, estimating the true expression based on the estimated measurement (imputation) errors and regressing phenotype on the true expression.

### 3.2 Simulation study

We used real genotype data to jointly simulate gene expression at both causal and non-causal genes in simulated reference and GWAS cohorts and continuous phenotypes in the GWAS cohorts, as done in prior work [16]. To calculate the recall and the effective false discovery rate (FDR), we tested single-gene associations against a phenotype generated using all of the simulated genes (Methods). For each simulated data set, we computed the the area under the precision-recall curve (AUPRC) of each method. We compared the AUPRC rather than the area under the receiver operating characteristic (AUROC) curve because the AUROC is not appropriate when the proportion of positive and negative examples is not 0.5 [?]. We ranked the genes according to the association statistic computed by each method, then computed the cumulative precision and recall (based on the simulated ground truth) for each position in the ranked list (Figure S2).

We simulated a reference cohort of 300 individuals and a GWAS cohort of 5,000 individuals for which 40 genes were causal and 1,000 were non-causal, and found that Bay-TS outperformed all other methods across the entire range of cis-regulatory architectures (Figure 1a). Interestingly, neither MI nor RC improved performance over OLS-E, which is likely explained by the fact that in this setting a central assumption in measurement error theory is violated. Specifically, when modeling the error in imputation, the second-stage predictors (means of imputed expression) and their associated errors (variances of imputed expression) are correlated because they both depend on genotype.

**Figure 1:**
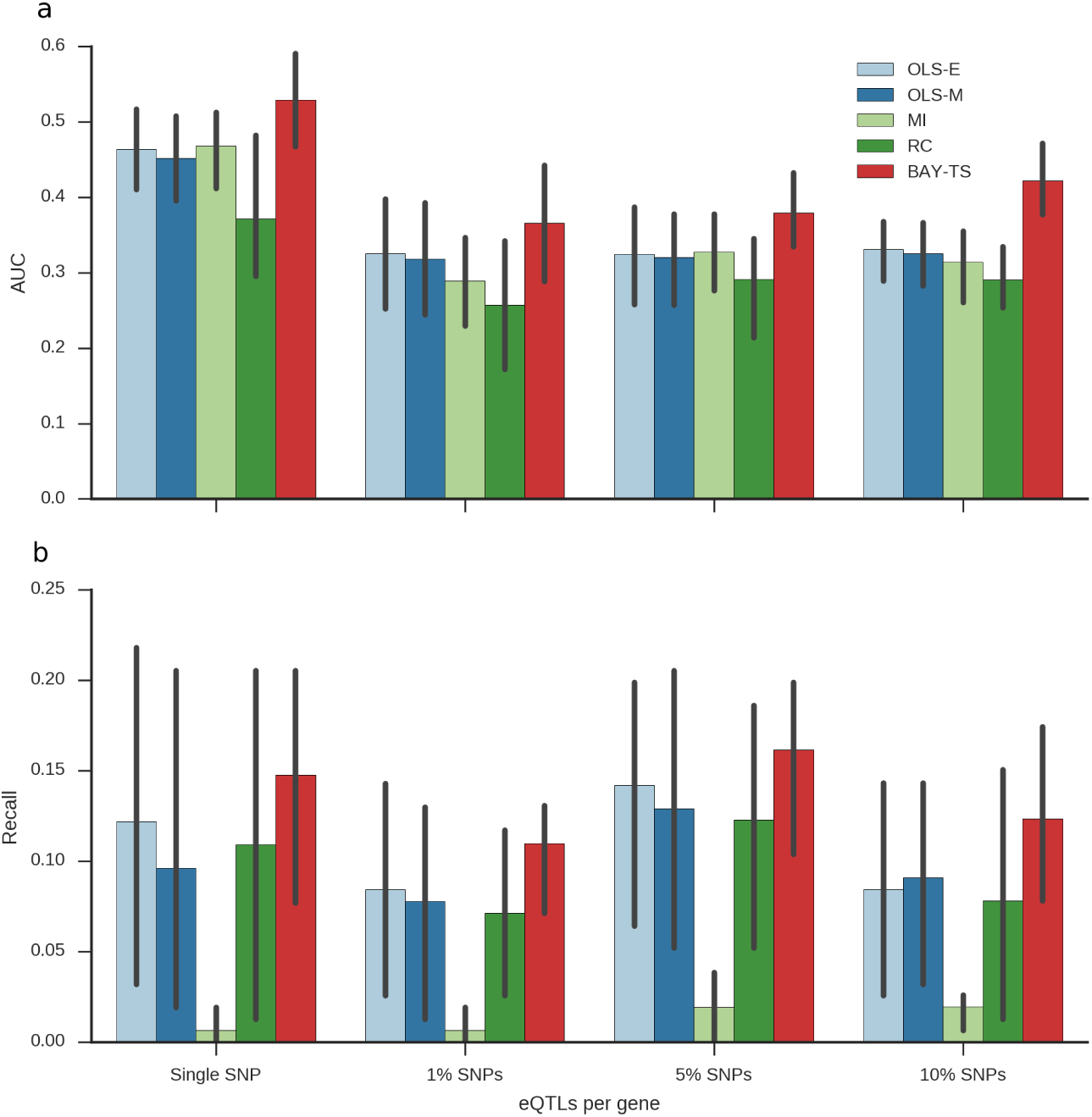
Simulation results. a) Area under precision-recall curve (AUPRC) for each method. Error bars represent standard error of AUPRC estimated over 4 replicates. b) Recall of each method controlling FDR at 10%. Error bars represent standard error of recall over 4 replicates.

We then investigated the recall of each method at FDR 10%. To control the FDR of BAY-TS, we calibrated a threshold for the Bayes factor (BF > 24) which controlled the average FDR over the entire range of simulation parameters at the desired level (Methods). For the other methods, we used the Benjamini-Hochberg procedure to control the FDR. We found that BAY-TS again consistently outperformed all the methods, with average recall equal to 15% (Figure 1b). Surprisingly, MI had the worst performance, likely due to deflated association statistics (Figure S3).

### 3.3 Application to seven diseases

Before applying our method to real data, we sought to evaluate the impact of expression normalization on the trained first-stage models in TWAS. We first asked which genes had gene expression reliably predictable from cis-genotypes. We used whole blood expression in 338 individuals from the GTEx project [4] and compared models trained on the published normalized expression values (GTEx-Norm) to models trained on regularized log transformed expression (RLog) (Methods). We found 1,666 and 1,655 genes with *R*^2^ ≥ 0 for GTEx-Norm and RLog, respectively, and found 1,043 genes common between the two sets (Figure 2a). However, only 987 of the 1,043 genes had enough non-missing data in the GWAS cohorts to successfully impute expression into GWAS.

**Figure 2:**
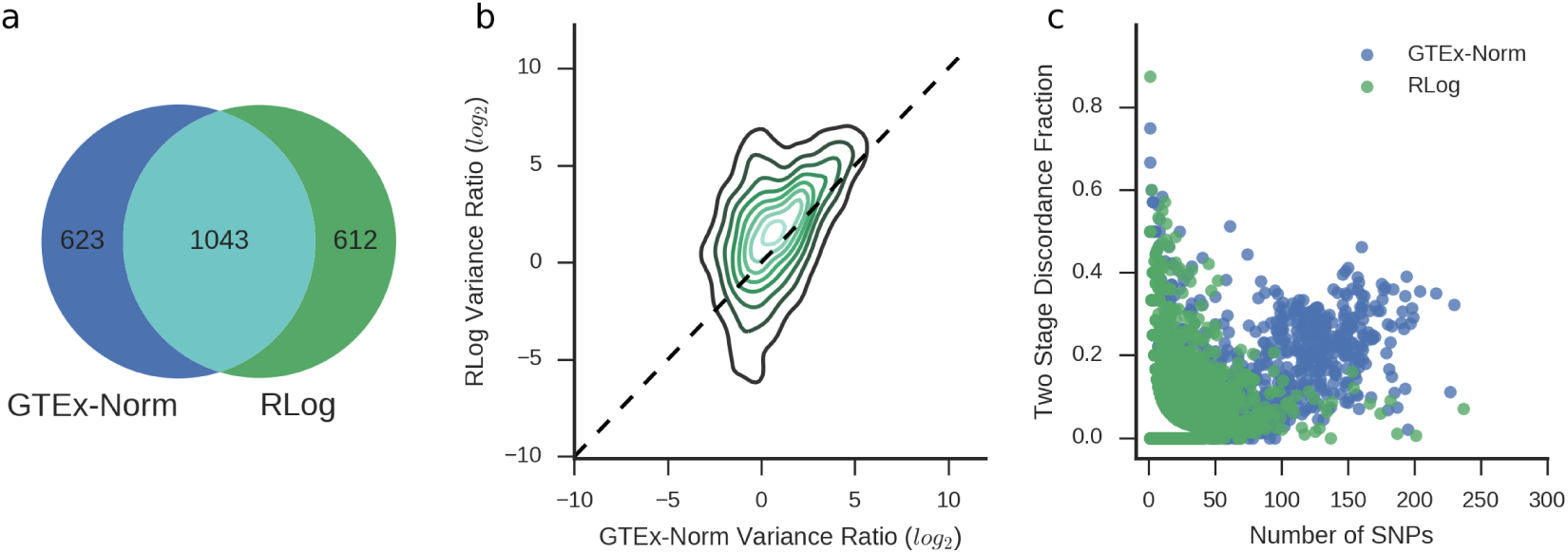
Impact of expression normalization on trained models of transcriptional regulation. a) The number of genes with cross-validation *R*^2^ ≥ 0. b) Density of the distribution of variance ratio (VR) for each of the 1043 common genes with cross-validation *R*^2^ ≥ 0. Contours denote surfaces of equal density. c) The total number of selected predictors per gene and the fraction of discordant predictors selected per gene.

We next considered the ratio between the variance of within-individual imputed expression estimates to across-individual estimates, which we define as the variance ratio (VR) (Methods). Measurement error theory predicts VR is correlated with power to detect associations, which we confirmed in simulation (Figure S4). Intuitively, after estimating the error variance of imputed expression, we can denoise the imputed expression by subtracting variation due to error in the imputed values. For the set of 987 genes, RLog based models had higher VR than GTEx-Norm based models (Figure 2b, S5), suggesting RLog normalization increases power to detect TWAS genes.

We next considered the ratio between the variance of within-individual imputed expression estimates to across-individual estimates, which we define as the variance ratio (VR) (Methods). Measurement error theory predicts VR is correlated with power to detect associations, which we confirmed in simulation (Figure S4). Intuitively, after estimating the error variance of imputed expression, we can denoise the imputed expression by subtracting variation due to error in the imputed values. For the set of 987 genes, RLog based models had higher VR than GTEx-Norm based models (Figure 2b, S5), suggesting RLog normalization increases power to detect TWAS genes.

We then asked whether the same cis-SNPs were reliably selected in the first stage model fitting for the 987 genes. We note that in this setting first-stage regression predictors are correlated due to linkage disequilibrium, and therefore regularized regression techniques such as elastic net will in general not select the same non-zero regression coefficients for replicate data sets. We calculated the discordance fraction between the predictors selected in our first-stage bootstrap training against the predictors included in a single model trained on the entire dataset (Methods). We found not only that GTEx-Norm models have more predictors per gene on average, but also have higher discordance in the selected predictors. Since RLog models find a more consistent set of cis-SNPs and have larger VR (Figure S5), we perform TWAS based on RLog models.

We performed TWAS on seven disease cohorts from the Wellcome Trust Case Control Consortium [9]: bipolar disorder (BD), Crohn’s disease (CD), coronary artery disease (CAD), hypertension (HT), rheumatoid arthritis (RA), Type 1 Diabetes (T1D), and Type 2 Diabetes (T1D). We first sought to replicate PrediXcan by using the published cis-regulatory model weights on our imputed genotypes [15]. We replicated only 18 of 40 reported TWAS genes (Bonferroni correction, *p* < 5.76 × 10^−6^), likely due to differences in imputation pipelines between the two studies. 13 of the discordant genes are in the Major Histocompatability Complex (MHC) region, and 15 have estimated effect sizes with the same sign as previously reported (Table S1).

We then performed TWAS using BAY-TS in each of the seven diseases and found 45 associated genes (FDR 10%, Table 1). Surprisingly, we only replicated 2 of 40 reported TWAS genes (Table S1). Moreover, we found that only 9 of the 40 reported genes in our set of high confidence genes with reliably predictable expression. The discrepancy in TWAS genes is due partly to differences in genotype imputation, expression normalization, and first stage model fitting as described above, and highlight the importance of data processing choices when performing TWAS. We compared BAY-TS to the other methods proposed above (including OLS-E, equivalent to PrediXcan [15]) and found that 23 of the 45 genes are not found by any of the other methods (Table S2, S3, S4). The other methods collectively did not find any associations for BD or CAD, likely due to lack of power.

**Table 1:** TWAS results for Bayesian two-stage regression model. BF: Bayes factor of BAY-TS. We only report genes with BF > 24, corresponding for FDR 10%. Effect size: Estimate of gene effect on disease liability. *R*^2^: cross-validation prediction accuracy. Ratio: variance ratio, the ratio of within-individual variance in imputed expression to across-individual variance. GWAS: best *p*-value in the WTCCC cohort within 1MB of the gene body. Catalog: best *p*-value reported in the NHGRI GWAS catalog within 1MB of the gene body. GEO: *p*-value for differential expression in independent case-control expression cohort

| Gene | Disease | Chr | BF | Effect size | R2 | Ratio | GWAS | Catalog | Diff. expr |
| --- | --- | --- | --- | --- | --- | --- | --- | --- | --- |
| SARDH | BD | 9 | 76.52 | -0.43 | 0.03 | 156.52 | 7.71e-04 | - | 0.98 |
| ZNF79 | BD | 9 | 25.18 | -1.01 | 0.04 | 23.03 | 1.40e-03 | - | 0.38 |
| WDR25 | BD | 14 | 28.07 | -0.69 | 0.15 | 41.10 | 6.76e-04 | - | - |
| PARL | CAD | 3 | 46.39 | 0.38 | 0.03 | 136.61 | 1.02e-04 | - | 0.26 |
| TMEM158 | CAD | 3 | 48.50 | -1.03 | 0.01 | 1.82 | 2.09e-03 | - | 0.092 |
| CYTH3 | CAD | 7 | 28.15 | -1.47 | 0.02 | 0.71 | 1.42e-04 | - | 0.47 |
| DAGLB | CAD | 7 | 77.12 | 2.13 | 0.09 | 4.13 | 1.42e-04 | - | 0.39 |
| FAAH | CD | 1 | 44.66 | 0.62 | 0.01 | 8.39 | 1.86e-04 | - | 0.38 |
| PIGC | CD | 1 | 311.50 | -1.73 | 0.12 | 34.47 | 5.93e-08 | 3e-22 | 0.0075 |
| HLA-B | CD | 6 | 9999.00 | -0.76 | 0.02 | 33.40 | 9.22e-09 | 7e-32 | 0.34 |
| HLA-DQA1 | CD | 6 | 10000.00 | 0.59 | 0.39 | 27.64 | 6.12e-08 | 9e-59 | 0.98 |
| PRKAB1 | CD | 12 | 26.17 | -0.87 | 0.03 | 74.76 | 1.33e-03 | - | 0.34 |
| LRRC37A2 | CD | 17 | 32.00 | -0.47 | 0.34 | 47.37 | 3.55e-05 | - | 0.086 |
| HSBP1L1 | CD | 18 | 29.86 | -0.88 | 0.25 | 32.46 | 3.05e-05 | - | - |
| CIAO1 | HT | 2 | 49.00 | 1.07 | 0.03 | 48.08 | 6.04e-05 | - | - |
| SHMT1 | HT | 17 | 9999.00 | 0.47 | 0.15 | 45.07 | 1.48e-04 | - | - |
| HLA-B | RA | 6 | 311.50 | -1.65 | 0.02 | 16.04 | 9.54e-33 | 3e-10 | 0.45 |
| HLA-DQA1 | RA | 6 | 10000.00 | 3.90 | 0.39 | 8.21 | 2.25e-86 | 1e-299 | - |
| HLA-DQA2 | RA | 6 | 10000.00 | 1.00 | 0.37 | 16.46 | 2.25e-86 | 1e-299 | 0.29 |
| HLA-DQB1 | RA | 6 | 10000.00 | 4.97 | 0.61 | 6.58 | 2.25e-86 | 1e-299 | 0.85 |
| HLA-DQB2 | RA | 6 | 10000.00 | 3.16 | 0.37 | 12.05 | 2.25e-86 | 1e-299 | - |
| HLA-DRB1 | RA | 6 | 10000.00 | -1.20 | 0.26 | 15.60 | 2.25e-86 | 1e-299 | 0.33 |
| HLA-DRB5 | RA | 6 | 10000.00 | -3.14 | 0.62 | 8.28 | 2.25e-86 | 1e-299 | 0.26 |
| HLA-G | RA | 6 | 4999.00 | -0.30 | 0.34 | 24.16 | 4.51e-16 | - | 0.75 |
| IER3 | RA | 6 | 10000.00 | 1.55 | 0.05 | 16.03 | 9.13e-12 | - | 0.0089 |
| MICA | RA | 6 | 10000.00 | 0.94 | 0.29 | 62.73 | 9.54e-33 | 3e-10 | 0.27 |
| BAK1 | T1D | 6 | 10000.00 | -1.10 | 0.14 | 32.53 | 2.35e-31 | - | 0.21 |
| BTN3A2 | T1D | 6 | 10000.00 | -0.47 | 0.19 | 148.71 | 1.94e-13 | - | 0.44 |
| HIST1H2AG | T1D | 6 | 10000.00 | -3.33 | 0.02 | 3.26 | 2.26e-15 | - | - |
| HLA-B | T1D | 6 | 10000.00 | -4.84 | 0.02 | 2.24 | 1.44e-91 | - | 0.37 |
| HLA-DQA1 | T1D | 6 | 10000.00 | 5.82 | 0.39 | 2.23 | 0.00e+00 | 5e-134 | 0.045 |
| HLA-DQA2 | T1D | 6 | 10000.00 | 9.79 | 0.37 | 5.93 | 0.00e+00 | 5e-134 | - |
| HLA-DQB1 | T1D | 6 | 10000.00 | 8.49 | 0.61 | 1.34 | 0.00e+00 | 5e-134 | 0.00018 |
| HLA-DQB2 | T1D | 6 | 10000.00 | 7.40 | 0.37 | 26.58 | 0.00e+00 | 5e-134 | 0.11 |
| HLA-DRB1 | T1D | 6 | 10000.00 | -3.97 | 0.26 | 8.23 | 0.00e+00 | 5e-134 | 0.24 |
| HLA-DRB5 | T1D | 6 | 554.56 | -2.08 | 0.62 | 0.46 | 0.00e+00 | 5e-134 | - |
| HMGN4 | T1D | 6 | 10000.00 | -1.44 | 0.09 | 9.59 | 1.94e-13 | - | 0.22 |
| IER3 | T1D | 6 | 10000.00 | 5.08 | 0.05 | 15.09 | 1.16e-48 | - | 0.13 |
| RBMS2 | T1D | 12 | 10000.00 | -3.31 | 0.23 | 1.23 | 8.93e-12 | - | 0.026 |
| RPS26 | T1D | 12 | 10000.00 | -3.10 | 0.02 | 12.37 | 8.93e-12 | 2e-25 | - |
| C1QTNF6 | T1D | 22 | 27.82 | 0.83 | 0.06 | 45.02 | 1.17e-05 | 2e-08 | 0.73 |
| PIGC | T2D | 1 | 33.01 | -0.98 | 0.12 | 30.33 | 1.73e-05 | - | - |
| KHK | T2D | 2 | 28.15 | -0.60 | 0.04 | 36.63 | 8.13e-04 | 2e-06 | 0.087 |
| TMEM158 | T2D | 3 | 103.17 | -1.06 | 0.01 | 1.77 | 9.53e-04 | - | 0.2 |
| POLR2J3 | T2D | 7 | 624.00 | -0.40 | 0.08 | 4.27 | 1.13e-03 | - | - |

We discovered three genes significantly associated with BD: SARDH, ZNF79, and WDR25. A higher burden of CNVs in the SARDH gene has been previously linked to BD and Schizophrenia[27], and its loss is the principal cause for sarcosinemia, which often leads to mental impairment[41]. However, ZNF79 and WDR25 are both poorly characterized and have not been studied in the context of BD previously.

We discovered four genes associated with CAD: DAGLB, CYTH3, PARL, and TMEM158. DAGLB is active in the triglyceride lipase activity pathway, and has been linked previously to high-density lipoprotein levels levels[51]. CYTH3 is a regulator of PI-3 kinase signaling, which mediates many pathways in the cardiovascular system [43]. Variants in PARL have been associated with increased levels of plasma insulin and predisposition to CAD [37]. In Chinese populations, the gene was also linked to higher levels of triglyceride and total cholestrol in both T2D and control populations [31]. Lower TMEM158 expression is associated with increased risk in both of the CAD and T2D populations in our study. It has previously transcriptionally associated with T1D, T2D, and gestational diabetes based on a case-control differential expression study performed using peripheral lymphomononuclear cells[11].

We discovered seven genes associated with CD (Figure 3): PIGC (1q24.33), FAAH (1p33), HLA-DQA1 (6pq21.32), HLA-B (6p21.33), PRKAB1 (12q24.23), LRRC37A2 (17q21.31), and HSBP1L1 (18q23). PIGC has been associated with inflammatory bowel disease as part of a larger Immunochip meta-analysis[24]. It is also significantly associated with T2D in our study. There has been no direct genetic evidence supporting the role of FAAH in CD, but recent experiments have shown that drugs targeting FAAH are effective against mouse models of colitis [40]. The HLA region has a known role in autoimmune disorders such as CD [13, 2]. PRKAB1 encodes the noncatalytic beta subunit of the AMP-activated protein kinase (AMPK), which has been previously experimentally validated to play an important role in IBD and is a therapeutic target for drugs used to treat CD and T2D [28]. Neither LRRC37A2 nor HSBP1L1 have been characterized in CD.

**Figure 3:**
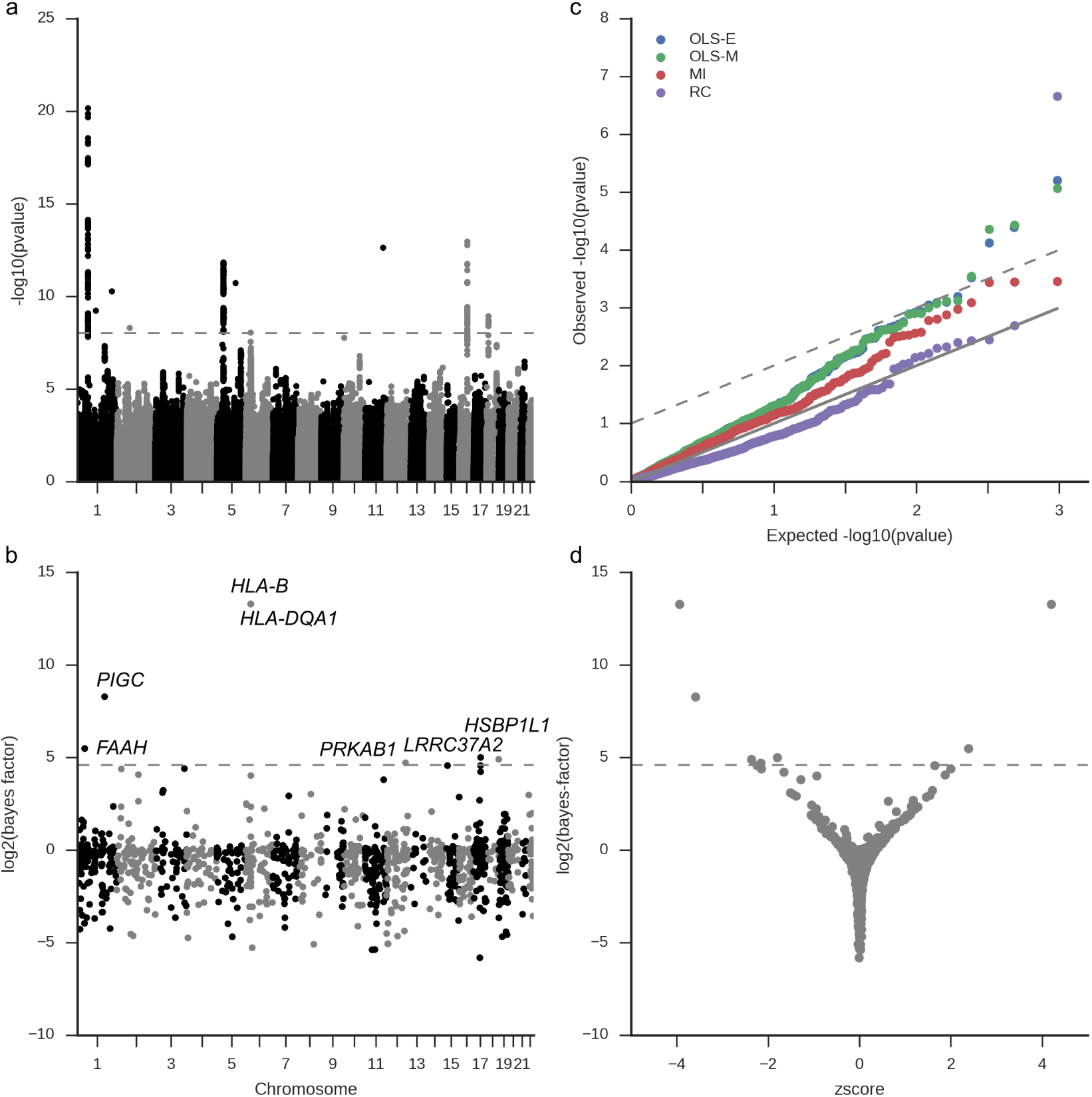
TWAS results for Crohn’s Disease. a) Manhattan plot of GWAS summary statistics estimated using FastLMM and LEAP on the WTCCC GWAS cohort. b) BAY-TS TWAS on the same samples reveals 7 genes that are significantly associated with Crohn’s Disease. c) Quantile-quantile plot of TWAS test statistics for frequentist methods. d) Bayes factor and z-score of BAY-TS associations. A Bayes factor of at least 24 corresponds to FDR 10%.

We found two genes associated with HT: SHMT1 and CIAO1. SHMT1 has previously been associated with hypertension, is a general prognostic marker for hypertension [34], is a marker for intra-cranial hypertension during space flight [45], and mediates the response to the angiogenesis inhibitor Bevacizumab[7]. CIAO1 encodes a protein in the iron-sulfur protein assembly complex and modulates activity of WT1, an oncogone associated with nephroblastomas [23]. It has not been studied in relation to HT.

We found multiple associations for RA in the MHC region: HLA-B, HLA-DQA1, HLA-DQA2, HLA-DQB1, HLA-DQB2, HLA-DRB1, HLA-DRB5, HLA-G, IER3, and MICA. Similarly, we found multiple MHC associations with T1D: BAK1, BTN3A2, HISTH1H2AG, HLA-B, HLA-DQA1, HLA-DQA2, HLA-DQB1, HLA-DQB2, HLA-DRB1, HLA-DRB5, HMGN4, and IER3. However, we discovered T1D associations with C1QTNF6, RBMS2, and RSP26. Larger meta-analysis of T1D has found association in the C1QTNF6 and RSP26 loci[12, 5]. However, RBSM2 has not been previously linked to T1D.

Finally, we found four genes are associated with T2D: KHK, PIGC, POLR2J3, and TMEM158. Ketohexokinase (KHK) plays a role in fructose metabolism and has been studied extensively as a possible cause for T2D [10, 26]. A sub-threshold association in its locus has been identified in a recent study of T2D in a Japanese population [21]. The PIGC locus has been previously linked to BMI[39], but not to T2D. POLR2J3 also has not been studied in T2D previously.

We further characterized our TWAS associations by looking at overlaps with GWAS loci (Methods). For each significant gene, we looked for a GWAS hit within 1 MB of the gene body (*p* < 5 × 10^−8^), and found overlaps for 23 of 42 genes. We then asked how many of the 42 genes were later discovered by larger meta-analyses in the seven diseases using the NHGRI GWAS Catalog [18, 50] and found 10 TWAS genes overlap loci reported in the catalog. However, most of the common genes found by both TWAS and GWAS lie in the MHC region. We finally sought to validate our TWAS associations using orthogonal case-control expression data sets[22, 6, 52, 8, 48, 25].

Surprisingly, we found only 3 TWAS genes are differentially expressed between cases and controls (limma modified t-test[46], Benjamini-Hochberg FDR < 0.1). We investigated the ranking of the top 250 genes by TWAS and differential expression and found no significant overlap between the ranked lists (Methods). There are several possible explanations for this discrepancy, including the tissue in which expression was measured, sample size, and technical confounders.

## 4 Discussion

Transcriptome-wide association studies (TWAS) have proven to be a powerful approach for identifying new genes associated with a phenotype by cleverly combining reference expression data and available GWAS data. TWAS directly associate genes to disease, revealing new biological insights. Indeed, these ideas have been extended to additional levels of mediation, incorporating histone modification data to study genetic effects on epigenomic control of transcription [17]. However, the field has not yet fully appreciated the importance of uncertainty in these multi-stage regression models. We showed that the state-of-the-art methods for TWAS adequately control type I error, but lose power due to uncertainty. We proposed a novel two-stage Bayesian method, BAY-TS, which outperforms not only existing methods but also standard methods from both missing data and measurement error theory. In applications to seven diseases from WTCCC, our method identified new genes not identified by previous methods, which do not incorporate uncertainty.

Our results reveal that uncertainty arises from many sources, not only differences in the trained models of regulation. Using the same GWAS genotypes, different imputation pipelines did not yield the same gene associations. We showed that expression normalization has an impact on trained models used for TWAS, and demonstrated that models trained on regularized-log transformed data were better than those trained on published GTEx expression data. Other studies have shown inconsistency between different population of reference transcriptomes from the same tissue [16]. These differences can be attributed to technical variation, environmental noise, sequencing technology, processing pipelines, and population differences. There is a pressing need to develop methods which adequately account for all these sources of uncertainty.

## 5 Methods

### 5.1 Uncertainty in TWAS

We assume a continuous phenotype *y*_*i*_ with zero mean collected on *n* individuals, and regress phenotype on predicted expression *w*_*i*_ for each gene whose expression levels can be predicted from genotype information. To handle binary phenotypes, we estimate latent liabilities using LEAP[49] and regress predicted expression against individual liabilities. For ease of exposition, we describe a model with no additional covariates; these can be included as additional terms in the model with no modification to the algorithms.

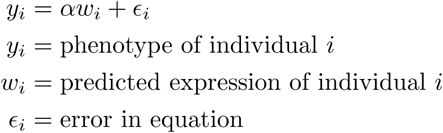

Current models use only one prediction for *w*_*i*_, based on a model of transcriptional regulation learned for the gene.

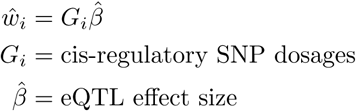

Here, we investigate models which account for the distribution of *w*_*i*_ and *β*. Assuming access to only one training cohort, we estimate these distributions by fitting *k* bootstrapped models regressing observed gene expression *E* on genotype *G*. Here, the regressions are performed using elastic net with regularization penalty tuned using cross-validation.

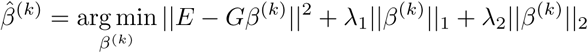

We estimate the first two moments of the distributions of *w*_*i*_ and *β* using the following equations:

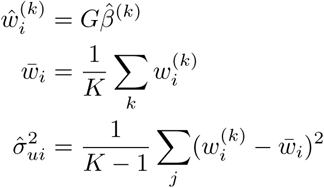

### 5.2 Bayesian two-stage regression model

To fit the Bayesian two-stage regression model for a gene (BAY-TS, Figure S1), we use the distributions of *β*_*j*_ learned using the *k* bootstrapped models for that gene as the prior in the second-stage regression.

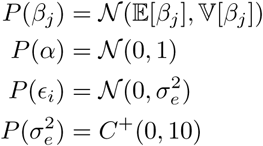

Here, 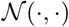 denotes the Gaussian density and 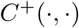 denotes the half-Cauchy density. Our inference goal is to estimate a Bayes factor comparing the model described above to a null model where *α* = 0. Rather than estimating the intractable model evidences and taking a ratio, we added a model indicator variable *z* and combined the null model 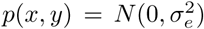 and the alternate model (described above). Then, the Bayes factor is given by the ratio *p*(*z* = 1)/*p*(*z* = 0). We implemented the model using PyMC3 [38] and used the Metropolis-Hastings algorithm to perform inference. We ran the MCMC chain for 100,000 steps and used the last 10,000 samples to compute the Bayes factor.

### 5.3 Multiple imputation

To perform multiple imputation (MI), we fit the *k* bootstrapped models for *w* against the phenotype using linear regression, calculate an aggregated test statistic *θ*_*mi*_, and compute a Wald test statistic.

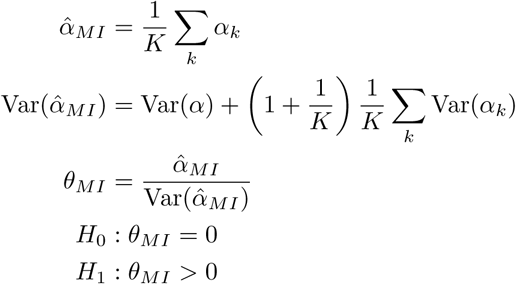

### 5.4 Regression calibration

We assume additive measurement error on the predicted expression value:

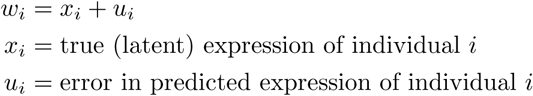

We assume measurement errors have zero mean and finite variance:

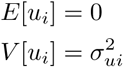

To perform regression calibration (RC), we impute the true expression value and regress phenotype against this estimated true expression. Given 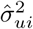, we regress *y_i_* on 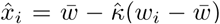, yielding estimate 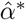. To estimate the association p-value, we perform a Wald test. We estimate Σ_*α*_, the covariance of 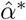 using a robust estimator [14]:

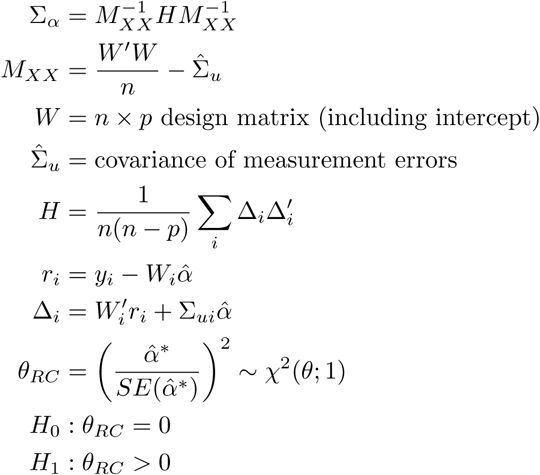

### 5.5 Simulation study

We used imputed dosages for 4,884 samples from the Hypertension, 58C, and NBS cohorts as described below. We selected 193 genes with cis-heritable gene expression (likelihood ratio test, GREML) in all of three studies as previously reported[16]: Metabolic Syndrome in Men (METSIM), Netherlands Twin Registry (NTR), and Young Finns Study (YFS). We held out 350 individuals as the training cohort and used the rest as the test cohort.

For each gene, we sampled the causal fraction of eQTLs from (Single, 1%, 5%, 10%) of SNPs from the cis-regulatory window. We computed the genetic value of each individual *X = Gβ* and added i.i.d. Gaussian noise to achieve proportion of variance explained (PVE) equal to 0.17 in expectation by sampling from 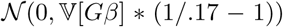, where 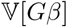 is the sample variance of the genetic values.

For each simulation, we sampled 40 causal genes and added i.i.d Gaussian noise to achieve PVE = 0.2 using the procedure described above. We computed the genetic value of each individual as *y* = *Xα* and add Gaussian noise as described above. We evaluated the method using sample sizes of 5000 individuals. We also tested the performance varying the number of non-causal genes from 400 to 4000.

### 5.6 GWAS processing

We downloaded Affymetrix genotypes for the Wellcome Trust Case Control Consortium seven diseases study in OXSTATS format called using the Chiamo algorithm from the European Genome Archive. We downloaded probe identifiers, hg19 positions, and strand information (http://www.well.ox.ac.uk/~wrayner/strand/) to convert positions to hg19 and used GTOOL version 0.7.5 to align all genotypes. We used PLINK version 1.09b to produce hard genotype calls with genotype probability threshold 0.99 and remove all SNPs and samples excluded from the original study.

We used SHAPEIT2 v2.r644 (ref. [19]) to exclude unalignable SNPs and phase the case and control cohorts independently for each autosome. We used default values for all model parameters. We used IMPUTE2 version 2.3.0 (ref. [20]) to impute into all SNPs and indels with MAF in European samples > 0.01. We divided the autosomes into 5 MB windows and threw out windows with fewer than 100 array probes.

For each cohort, we hard-called imputed dosages with genotype probability threshold 0.9. For each disease cohort, we produced a case-control set of hard-called genotypes for both the array genotypes and imputed genotypes by merging all chromosomes with the shared controls (1958 Birth Cohort and National Blood Services). We used GCTA 1.24 (ref. [53]) to estimate a genetic relatedness matrix on the case-control array genotypes and prune pairs of individuals with relatedness > 0.05. We used plink to remove these individuals from the imputed genotypes, and further remove indels and SNPs with missingness > 0.01, differential missingness (*p* < 0.05) or HWE *p* < 10^−5^.

We used LEAP version 0.1.8.9 (ref. [49]) to estimate latent liabilities for each chromosome of each case-control dataset separately (holding out that chromosome) using the array genotypes. We used FastLMM version 0.2.26 (ref. [29]) to compute association *p*-values for the imputed genotypes using kinship matrices estimated from the array genotypes (described above). We made extensive use of GNU parallel[47] to facilitate the analysis.

### 5.7 Reference expression processing

We downloaded genotypes and RNA-Seq read counts in the v6 release of GTEx from dbGaP. We restricted our analysis to only those genes which had RNASeqC gene-level read count >= 10 in at least 10 individuals, resulting in 12,049 genes. We transformed the counts to regularized log-transform values (RLog) using DESEq2, adjusting for sequencing depth using the median-of-ratios method [32]. We trained our models on these normalized values to models trained on the published expression values (GTEX-Norm).

We extracted genotypes for SNPs within 500kb upstream and downstream of the transcription start and end sites for each gene using plink. We filtered sites with missingness > 0.01 or HWE *p* < 10^−5^. For models fit on the published expression values (GTEx-Norm), we included 3 genotype principal components (PCs), 35 expression PEER factors, gender, and sequencing platform as covariates. For models fit on RLog, we used the same covariates but used 10 expression PCs instead of the PEER factors.

To find the optimal elastic net penalty parameter and l1/l2 regularization ratio, we used Elastic-NetCV from the scikit-learn package [36] with possible l1 ratio values 0.1, 0.2, 0.3, 0.4, 0.5, 0.6, 0.7, 0.8, 0.95, and 0.99 and 100 uniformly distributed penalty parameters from 0.1 to 1. Using the best fitted penalty parameters parameters, we fit 50 bootstrapped models by sampling 300 individuals from the 338 samples.

We imputed expression using PrediXcan software as well as an independent implementation. In our independent implementation, we filtered all GWAS sites that had higher than 10% missing genotypes. Additionally, we assigned the average value to all missing genotypes (which is possible for best-guess imputed genotypes after thresholding on the posterior probability of any genotype call). We note that the implementation of PrediXcan assumes there is no missingness in the data.

To calculate cross-validation prediction accuracy, we predicted hold-out gene expression using only genotype (omitting covariates). We used 5-fold cross-validation and calculated the average *R*^2^ across the folds.

We define the variance ratio (VR) as the ratio of variance of mean imputed expression to the mean of the variance of imputed expression. Intuitively, the VR compares the within-individual variation in imputed expression to between-individual variation. We estimate VR by estimating the necessary means and variances over the 50 bootstrapped models described above.

### 5.8 GEO data

We downloaded case-control expression datasets for the seven diseases from GEO and processed them using the GEO2R web service: GSE12654 (BD) [22], GSE20681 (CAD) [6], GSE6731 (CD) [52], GSE703 (HT) [8], GSE15573 (RA) [48], GSE55100 (T1D) [54], GSE21321 (T2D) [25]. For each data set, we found differentially expressed genes between cases and controls using limma and the GEO2R web service. We assessed significance in the ranking between TWAS lists and GEO lists using the R package OrderedList [55], restricting to the top 250 overlapping genes for each disease.

